# Bringing Diffuse X-ray Scattering Into Focus

**DOI:** 10.1101/219113

**Authors:** Michael E. Wall, Alexander M. Wolff, James S. Fraser

## Abstract

X-ray crystallography is experiencing a renaissance as a method for probing the protein conformational ensemble. The inherent limitations of Bragg analysis, however, which only reveals the mean structure, have given way to a surge in interest in diffuse scattering, which is caused by structure variations. Diffuse scattering is present in all macromolecular crystallography experiments. Recent studies are shedding light on the origins of diffuse scattering in protein crystallography, and provide clues for leveraging diffuse scattering to model protein motions with atomic detail.

## Introduction

With over 100,000 X-ray structures deposited in the wwPDB [1], improvements in data processing pipelines, and the advent of completely unattended data collection, it seems hard to imagine that there are any aspects of protein X-ray crystallography that remain to be optimized. However, only about half of the X-rays scattered by the crystalline sample are currently being analyzed – those in the Bragg peaks. The weaker, more smoothly varying features in diffraction images, known as diffuse scattering, are largely ignored by current practices. While the analysis of diffuse scattering is an established method in the fields of small molecule crystallography [2] and materials science [3], there are only very few foundational studies of diffuse scattering in macromolecular crystallography [4-18]. However, the relative scarcity of diffuse scattering studies is poised to change as activity in the field has recently increased.

A small group of researchers (including MEW and JSF) met in 2014 to discuss the challenges and opportunities of investigating macromolecular diffuse scattering [22]. Our attention was drawn to several key developments in the field of macromolecular crystallography that motivated and enabled assessment of the diffuse signal. First, structural models obtained using traditional methods appear to be reaching a plateau in quality, as R factors remain relatively high compared to what can be achieved in small-molecule crystallography. The origin of this “R-factor gap” is likely due to the underlying inadequacies of the structural models refined against crystallographic data [23]. These inadequacies can only be overcome if we can improve the modeling, including, for example, conformational heterogeneity (especially in data collected at room temperature [24]), solvation, and lattice imperfections that break the assumptions of “perfect crystals” used in data reduction and refinement. Second, new detectors were enabling collection of data with lower noise, higher dynamic range, and highly localized signal. Third, new light sources were emerging with very bright, micro-focused beams (e.g. X-ray free-electron lasers). Collectively, these factors made us optimistic that diffuse scattering data both was needed and could be measured accurately enough to improve structural modeling. In early 2017, many of us met again to discuss the progress of the field with respect to each of these challenges identified in 2014 [25]. In this review, we provide our perspective on this progress and the status of the field, informed in part by our observations at that meeting and advances covered by Meisburger et al. [26]. While there have been exciting developments in recent years, there are still major challenges ahead, include modeling atomic motions in protein crystals using diffuse scattering data with accuracy comparable to the Bragg analysis, and utilizing these models of protein motions to distinguish between competing biochemical mechanisms.

## Data collection

Extraction of diffuse scattering data from conventional protein crystallography experiments is becoming straightforward thanks to the increased accessibility of photon-counting pixel array detectors (PADs, e.g. Pilatus detectors). These detectors have greater dynamic range and do not suffer from “blooming” overloads that obscured diffuse signals near Bragg peaks on conventional charge-coupled device (CCD) detectors. (An early CCD detector was programmed to drain excess charge away from overflowing pixels to enable measurement of diffuse scattering data [17,27]; however, this feature was not implemented in commercial detectors.) Additionally the use of PADs has led to changes in data collection strategies, such as the use of fine phi angle scans, that facilitate analysis of Bragg peaks and diffuse features from the same set of images [19]. A second major advance is the measurement of diffuse scattering using an X-ray free-electron laser (XFEL) in a serial femtosecond crystallography (SFX) experiment [28]. Using an XFEL enables collection of radiation-damage-free room temperature data, as well the potential to examine time-resolved changes in the diffuse scattering signal.

Despite these advances in collection of diffuse scattering data, minimizing background scattering remains the most important obstacle to collecting high quality data. While it is possible to remove some background scattering during data processing, the cleanest separation requires one to remove scattering extraneous to the crystal during the experiment. Factors to consider during collection of single crystal datasets include the thickness and orientation of the loop (for relevant mounting schemes), the volume of liquid surrounding the crystal, and the amount of airspace between the crystal and the detector. Background air scatter can be also reduced by a Helium or vacuum path between sample and detector. Collection of SFX data adds additional complexity, as the injection stream and crystal size will vary. Ayyer et al [28] addressed this challenge by selecting only the frames with the strongest diffuse scattering signal, in which the size of the crystal was expected to be comparable to the width of the jet. As the landscape of sample delivery devices for SFX and conventional crystallography continues to evolve, mounted sample delivery on materials such as graphene [29] provides a promising route for minimization of background scattering.

## Data integration

Early studies of protein diffuse scattering focused on explaining features in individual diffraction images. The introduction of methods for three-dimensional diffuse data integration enabled quantitative validation of models of correlated motions [17]. Several approaches to 3D data integration now have been implemented [27,28,30-32]. These approaches differ in several key ways: (1) the scaling of intensities when merging the data; (2) the handling of intensities in the neighborhood of the Bragg peak; and (3) the strategy for sampling of reciprocal space. In the *Lunus* software for diffuse scattering (https://github.com/mewall/lunus) we have chosen:

1. To use the diffuse intensity itself to scale the diffuse data (as opposed to using the Bragg peaks, as in Ref. [31]). This choice avoids artifacts due to potential differences in the way the Bragg and diffuse scattering vary with radiation damage and other confounding factors. The response of these signals to damage requires further study before a definitive scaling strategy can be chosen.
2. To ignore or filter intensity values in regions where the variations are sharper than the 3D grid that will hold the integrated data. This can include masking halo intensities too close to a Bragg peak, and kernel-based image processing to remove Bragg peaks from diffraction images. These steps avoid the mixing of signal associated with sharp features into the signal associated with larger-scale, cloudy diffuse features. The sharply varying features (e.g. streaks) are an important component of the signal; however, to avoid artifacts in analysis, we prefer to measure them on a grid that is fine enough to resolve them [17]. If the sampling is finer than one measurement per integer Miller index, but still too coarse to resolve the halos, and if the halo intensity is nevertheless included (as in Ref. [31]) then the measurements at integer Miller indices may be segregated from the rest of the data and analyzed separately.
3. To sample the data on a grid that includes points at Miller indices (corresponding to where the Bragg peaks are located), and, for finer sampling, points corresponding to integer subdivisions of Miller indices. Sampling strategies that are not tied to the reciprocal lattice also are valid (as used in Refs. [28,30]); however, on-lattice strategies enable leveraging of existing crystallographic analysis and modeling tools for diffuse scattering.

Efforts are now underway to decrease the burden of diffuse data integration and make diffuse data collection accessible for any protein crystallography lab. Recent algorithmic improvements have led to scalable, parallelized methods for real-time processing of single-crystal synchrotron data, decreasing the time required to extract a diffuse dataset from diffraction images. These improvements aim to keep pace with real-time analysis of Bragg data at high frame rates, such as those expected at LCLS-II and euXFEL. Initial tests mapped staphylococcal nuclease diffuse data onto a fine-grained reciprocal lattice, using two samples per Miller index [33]. This implementation of the *Lunus* software is capable of processing thousands of diffraction images within a few minutes on a small computing cluster.

In addition to improving the scalability of diffuse scattering data processing, efforts are underway to create a push-button diffuse data processing pipeline. The *Sematura* pipeline (https://github.com/fraser-lab/diffuse_scattering) was inspired by the user-friendly environment provided by software for analyzing Bragg peaks, such as xia2 [34]. To ensure portability the project was built upon the CCTBX framework [35], with future work focusing on developing *Sematura* as a CCTBX module for ease of access.

## Building and refining models of protein motions

### Liquid-like motions

After early experiments on tropomyosin [15], the liquid-like motions (LLM) model became a key tool in interpreting diffuse features in diffraction images [4,6]. In the LLM model, the crystal is treated as a soft material. All atoms are assumed to exhibit statistically identical normally distributed displacements about their mean position. The correlation between atom displacements is a decreasing function of the distance between the atoms, usually an exponential decay. A LLM was used to interpret early 3D diffuse data sets, refined using a correlation coefficient as a target function [17]. Successful refinement of a LLM model was used to demonstrate that interpretable diffuse datasets can be extracted from Bragg diffraction experiments, when data collection is not specifically targeted at measuring the diffuse signal [19].

Peck et al. [31] recently found the ability of the LLM to include correlations across unit cell boundaries was essential for modeling the diffuse signal in several 3D datasets. This result is intuitive, as many atoms in a typical protein crystal are within 5-10 Å of symmetry related molecules. Overall the LLM model has proven to be a simple means of explaining the general features of the data with a straightforward interpretation, and therefore remains an important first approach to analysis of protein diffuse X-ray scattering data.

### Normal mode analysis and elastic network models

Beyond the LLM model, normal mode analysis (NMA) of elastic network models (ENMs) can provide insights into the soft modes of protein dynamics in more detail, helping to reveal mechanisms that bridge protein structure and function [36]. In an ENM, the atoms of the crystal structure are connected by springs, and the resulting network is coupled to a thermal bath. NMA then yields the covariance matrix of atom displacements. The diagonal elements of the covariance matrix correspond to the crystallographic B factors. ENMs are often used to predict B factors, which come from the Bragg analysis through the crystal structure model. However, Riccardi et al. [37] showed how to renormalize the entire covariance matrix using the crystallographic B factors. Importantly, this renormalization enables any ENM to be entirely consistent with the Bragg data, while preserving differences in diffuse scattering. Diffuse scattering could help differentiate between these ENMs because the off-diagonal elements directly influence the diffuse signal. Thus, there is an opportunity for carefully measured diffuse data to be used in refinement of ENM models, and subsequent refinement of models of protein structure and dynamics.

Indeed, many key elements needed for refinement of normal modes models using diffuse scattering already have been demonstrated. Cloudy diffuse features in X-ray diffraction from lysozyme crystals resemble the diffuse scattering predicted from simulations of normal modes models [9,13]. Similarly, sharper diffuse features in the neighborhood of Bragg peaks in ribonuclease crystals can be captured by lattice normal modes [38] Different varieties of ENMs for staphylococcal nuclease give rise to distinct diffuse scattering patterns, even when renormalized using the crystallographic B factors [37].

Three-dimensional diffuse scattering data from trypsin and proline isomerase (CypA) recently were modeled using ENMs [19]. The agreement was substantial, considering that the models were not refined. On the other hand, Peck et al. [31] found a low agreement between ENM models and diffuse data. How much can refinement improve the agreement of an ENM model? Here we provide an example. In our example, the asymmetric unit of PDB ID 4WOR was expanded to the P1 unit cell, and an ENM was constructed as in Ref. [19]. The spring force constants between C-alpha atoms were computed as *e*^−*r_ij_/λ*^, where *r*_*ij*_ is the closest distance between atoms *i* and *j*, either in the same unit cell or in neighboring unit cells of the crystal structure. All atoms on the same residue as the C-alpha were assumed to move rigidly as a unit. The initial value *λ* = 10.5 Å yielded a linear correlation of 0.07 with the anisotropic component of the diffuse data, as computed in Ref. [19]. Powell minimization using the *scipy.optimize.minimize* method was used to refine the value of *λ*, using the negative correlation as a target. The final correlation was 0.54 for a value *λ* = 0.157 Å – a substantial improvement, but one that indicates that the direct interactions are essentially limited to nearest atomic neighbors. Simulated diffuse intensity in diffraction images calculated using the model vs. the data show similarities in cloudy diffuse features (Fig. 2). Key strategies for improving the model are: extending from a C-alpha network to an all-atom network; using crystalline normal modes that extend beyond a single unit cell (prior studies used the Born von Karman method to compute these modes [37,38], but did not fully include the resulting modes in the thermal diffuse scattering calculation [39]); and allowing spring constants to deviate locally from the exponential behavior. Optimizing this type of model has applications beyond diffuse scattering validation and model refinement, as structures derived from normal modes analysis of network models have been useful for providing alternative starting points for molecular replacement [40] and have recently been used in an exciting local refinement procedure in cryo electron microscopy [41].

**Fig. 2.**
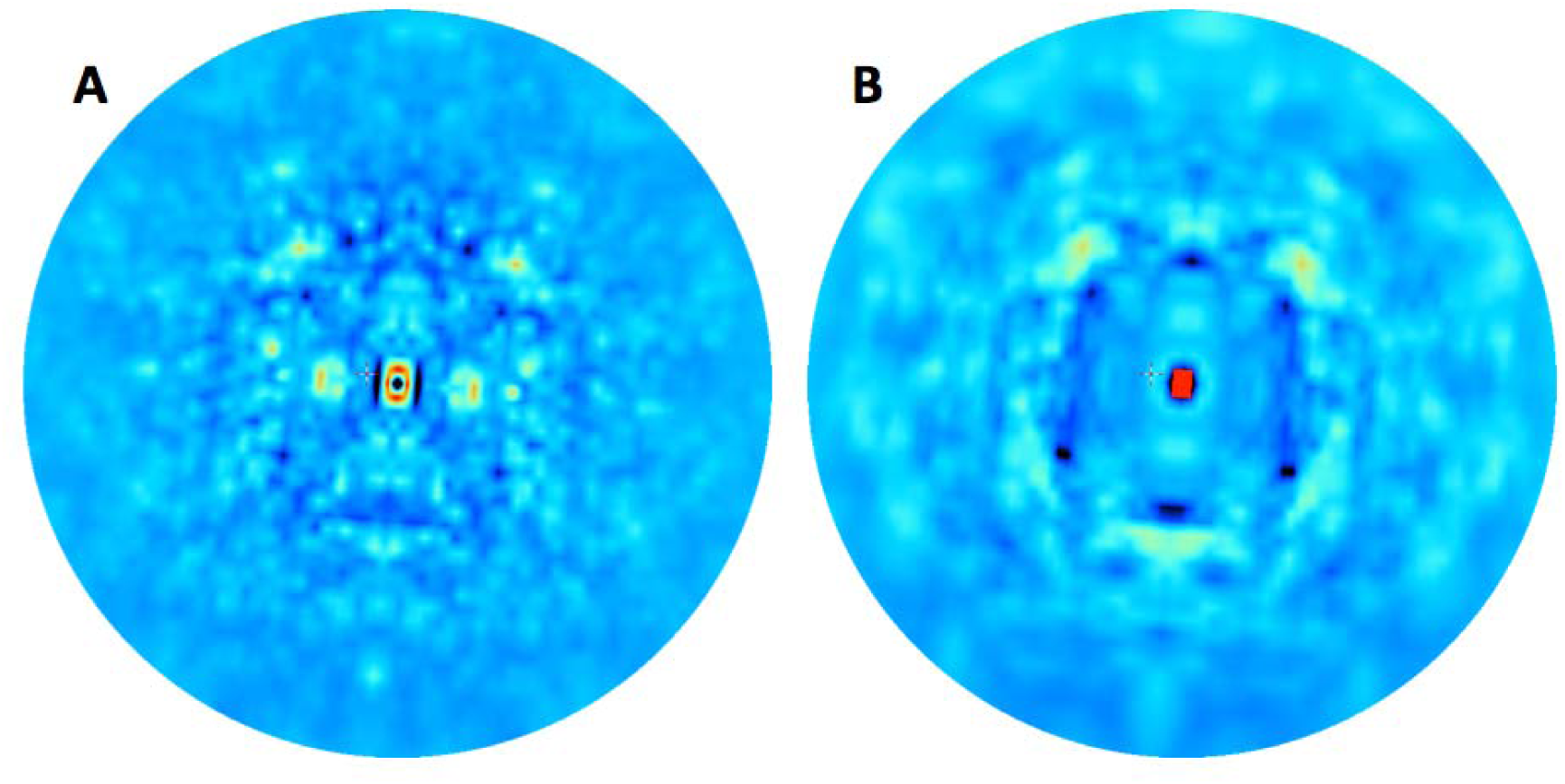
Comparison of simulated diffuse intensity in diffraction images computed from (A) a refined ENM of staphylococcal nuclease and (B) experimental data from Ref. [17].

### Ensemble refinement

A great promise of diffuse scattering is the potential to validate ensemble or multiconformer models of protein structures (Figure 1). As for as with Translation-Libration-Screw (TLS) refinement [42] and ENM models, diffuse signal might differ for ensembles that result in the same average structure. Even if information about atomistic conformations remains out of reach, the signal could potentially be leveraged to improve ensemble models derived from time-averaged refinement using the scheme by Gros and colleagues [43]. Currently, this procedure operates on the rationale that large scale deviations can be modeled using a TLS model, and the residual local deviations are then sampled by a molecular dynamics simulation with a time-averaged difference electron density term. Our work has revealed that diffuse scattering calculated from TLS models of disorder do not match the measured diffuse signal, however, indicating that TLS is a poor descriptor of the disorder within the protein crystals we considered [19]. Given the improvements seen when including neighboring unit cells in LLM models [31], the disorder of the crystal environment might be better accounted for by a coarsegrained model of intramolecular motion using a NMA model refined against the diffuse scattering signal. In addition, due to the limited number of copies in ensemble models, they can exhibit artificially long length scales compared to molecular dynamics simulations, which contain orders of magnitude more finely time-sliced “snapshots” of the system[44]. Ensemble models of diffuse scattering data will therefore need to include the effect of decoherence corresponding to the finer scale motions that are filtered out in conformational selection. Once large-scale disorder is accounted for by NMA, local anharmonic deviations from the modes can be explored using MD simulations restrained by the X-ray data. As diffuse analysis becomes more sensitive, the selection of the final representative ensemble also might be optimized against the diffuse data. This selection step could supplement the current practice of selecting an ensemble that matches the Bragg data.

**Fig. 1.**
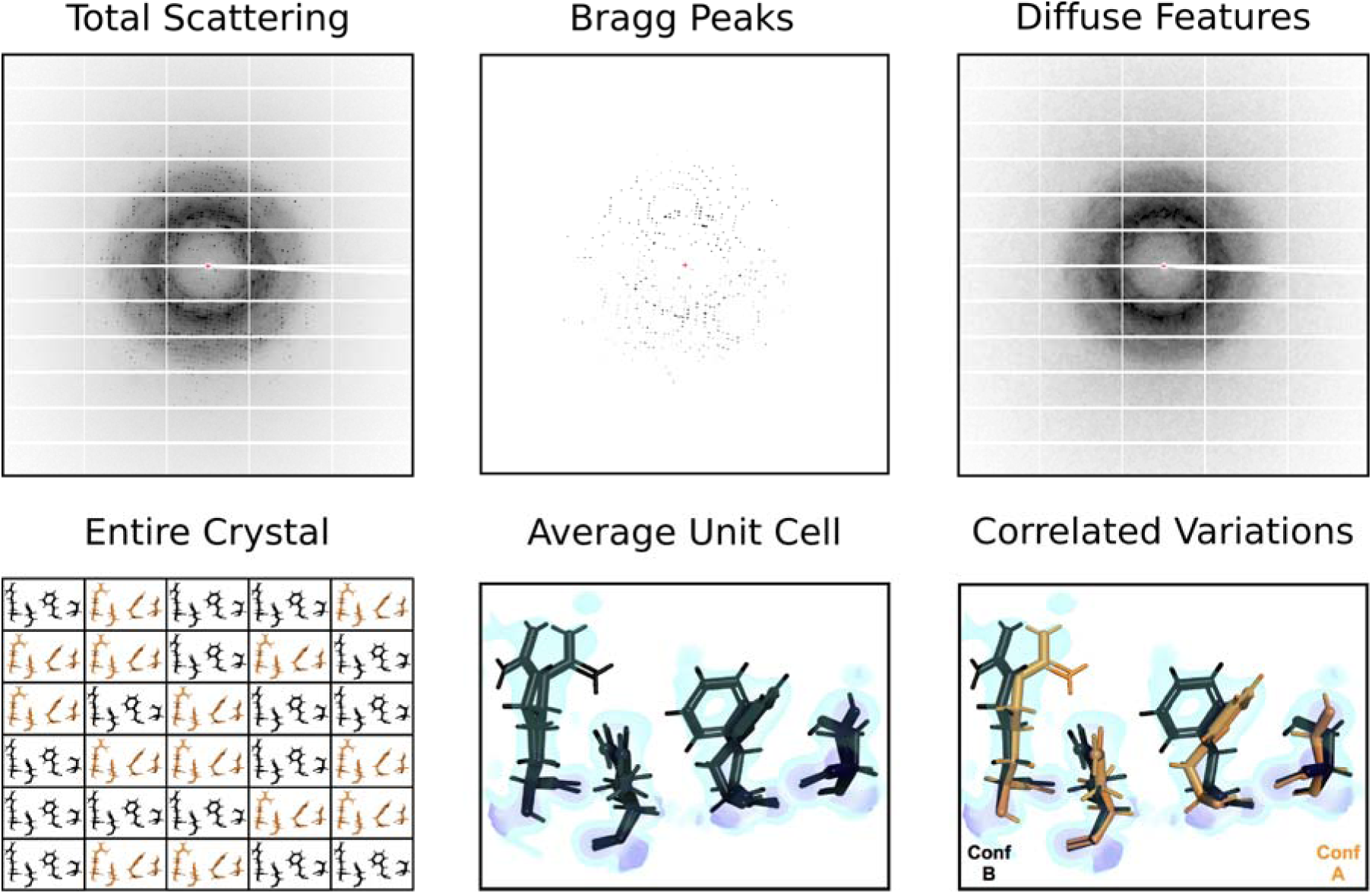
A typical detector image in X-ray crystallography (from [19]) (*upper, left*) records all of the X-rays scattered by a protein crystal during a single exposure. Dark pixels correspond to high X-ray intensities. A cartoon crystal is depicted (*lower, left*) that contains a series of unit cells, with the contents of any given unit cell adopting one of two conformations (the conformations are expected to be more varied in a real protein crystal). Conformation A is shown in orange, while conformation B is shown in black (*lower panel*). During analysis, data are reduced by examining only the Bragg peaks (*upper, middle*), which report on the average charge density within a unit cell (*lower, middle*). The electron density is shown in blue, with areas of especially strong charge highlighted in purple. While multiple conformations may be modelled into the average density, assigning which conformations occur together across residues requires additional information. Current modeling practices use geometric constraints to help classify different alternative conformation groups [20]. The diffuse scattering left behind during data reduction (*upper, right*) is an additional potential source of such information. Diffuse scattering includes an isotropic component that is determined both by protein and solvent scattering [21,22], and an anisotropic component that is dominated by correlated protein motions within the crystal [22]. Analyzing this anisotropic signal might help to distinguish networks of residues that mo ve together (*lower, right*).

## Molecular dynamics simulations

In addition to refining models of protein motions, diffuse scattering can be used to validate MD simulations [7,9,21,22,45-47]. Early efforts were hindered by the use of 10 ns or shorter simulation durations [7,9,21,45], which lacked sufficient sampling for the calculations. Microsecond duration simulations of protein crystals are now becoming routine [22,33,48,49]. For staphylococcal nuclease, microsecond simulations overcome the sampling limitations for diffuse scattering calculations, while providing insight into ligand binding and catalysis [22].

The agreement of the total diffuse intensity with MD simulations is high for staphylococcal nuclease [22,46], yielding a linear correlation of 0.94 for a microsecond simulation [22]. Agreement with the 10-fold weaker anisotropic component is lower [22,33], but is more sensitive to the details of the simulation, creating opportunities for increasing the accuracy of MD models. Expanding the staphylococcal nuclease model from a single periodic unit cell to a 2×2×2 supercell increased the correlation with the anisotropic component to 1.6 Å resolution from 0.42 to 0.68 for a microsecond simulation [33].

Even though MD simulations provide a picture of crystalline dynamics at atomic detail, the accuracy of the published MD models is not yet high enough to validate the atomic details using diffuse scattering. Although we do not know what level of accuracy will be required for diffuse scattering to reveal the atomic details of protein motions, the possibility of validating the atomic details of protein motions strongly motivates improving the MD models. Ideas for improving the MD model include: increasing the size of the supercell even further, to 3×3×3 or beyond; improving force fields; increasing the simulation duration; and introducing crystal imperfections such as vacancies (missing copies of the protein) or dislocations. It is also possible that higher quality experimental data would be required to improve the model. Additional insights for increasing model accuracy might come from solid state NMR (ssNMR) experiments combined with crystalline protein simulations [50-52], which create opportunities for joint validation of MD simulations using crystallography and NMR.

## Phasing and resolution extension

In a high-profile publication, the Chapman and Fromme groups integrated the first three-dimensional diffuse scattering dataset from a serial femtosecond protein crystallography experiment at an X-ray free electron laser [28]. Their analysis focused on the potential for phasing and resolution extension of a charge density map of photosystem II (PSII). In this study, the method, based on the difference-map algorithm [53], depends critically on the assumption that the diffuse signal is proportional to the molecular transform of the PSII dimer. In this respect, the work is closely related to that of Stroud and Agard [54] and Makowski [55] on phasing using continuous diffraction data. In Ref. [28], and in a follow-on study using improved data integration [56], the validity of the method was argued by assuming independent, rigid body motions of the dimer.

Using diffuse scattering for phasing and resolution extension might prove to be useful in rare cases when diffuse scattering extends to higher resolution than the Bragg diffraction; however, many technical questions remain both about the origin of diffuse scattering in PSII and the role of diffuse scattering in yielding the resulting charge density. What effect does the presence of Bragg peaks have on phasing and resolution extension in the 4.5-3.5 Å range, which is where the diffuse intensity was measured? How does the improvement in the PSII map compare to what would be obtained by using randomized intensities, due to the free lunch effect [57] and solvent flattening [58]? How would the R-factors in the extended resolution range reported in PDB 5E79 compare to pseudo-crystallographic refinement [59] of using either random intensities or the uniform average intensity in these bins? How robust are the improved features of the charge density in Ref. [28] to omit map analysis [60], especially at the solvent/protein interface? Might a LLM model (or a ENM or MD model) more accurately describe the diffuse scattering than rigid-body translations of PSII dimers (in Ref. [56], the agreement was improved when the intensities were convoluted with a 4×4×4 voxel kernel)? Can the model be improved by assuming the rigid units are coupled instead of independent [8], or if the model included rotations as well as translations [14] (in Ref. [56], the intensities were rotationally blurred, but this does not correspond to rigid-body rotations [61])? What is the role of substitution disorder [62] (e.g. unit cells in which one or more copies of the PSII dimer are missing) in determining the diffuse signal? Understanding the implications of this method for protein crystallography will rely on answering these questions and determining whether signals at higher angle than Bragg are commonly observed.

## Future perspective

The massive investment in structural genomics in the 2000s dramatically increased the robustness of X-ray crystallography data collection, processing, and refinement. The resulting technological improvements and standardizations have led to more robust methods and instrumentation for data collection that are well-aligned with the requirements for diffuse scattering experiments, enabling measurement of diffuse scattering data from traditional crystallography experiments [19,31]. These advances, along with software that makes the data processing and analysis more accessible, will enable diffuse scattering studies at any modern beamline, by any crystallography lab. Why should crystallographers take advantage of this offering? Diffuse features in protein crystallography can be myriad and complex: a mixture streaks, satellite reflections, isotropic scattering, cloudy patterns, circles, and others. Sometimes the diffuse signal appears to reflect imperfections in crystal packing, and offers a possible explanation for why a structure cannot be solved [18]. In other cases, the diffuse signal might be so complicated as to be uninterpretable, or might be so weak that it is difficult to learn anything interesting.

Integrating diffuse scattering with Bragg diffraction to improve crystallographic models could become a major application [17,32,63]. Although assuming proteins are rigid provides the greatest potential for phasing using diffuse scattering data [28,56], multiple studies of both Bragg and diffuse scattering point to a more dynamic picture of crystalline proteins. A model with internal motions such as the LLM tends to obscure the molecular transform signal and to limit the information to what is available from the crystal transform, at Miller indices [31]. Nevertheless, because the diffuse signal can extend well beyond the resolution limit of the Bragg peaks in rare cases, it still allows for the possibility of resolution extension. The blurring of the signal implied by the LLM means there is a loss of information in the diffuse Patterson function at long distances, so the path to resolution extension might require model refinement in addition to, or instead of, direct methods. In addition, the apparent success of the LLM [4,6,17,19,31] and MD simulations [21,22,33,46,47] in obtaining insights into diffuse scattering data points to a picture in which internal motions are important. This opens up the possibility that diffuse scattering can be used to reveal atomic models of protein motions, a possibility that is eliminated when proteins are treated as rigid units.

Ironically, the strongly diffracting model systems that have enabled experimental measurements of diffuse scattering may contain less informative signals than more poorly ordered crystals. In the case of poorly ordered systems, multiple significant conformational minima may co-exist in the crystal. This disorder would limit the power of Bragg diffraction, while the correlations present in the ensemble or the spread of conformations between extremes [64] would lead to diffuse signal. However, increased disorder, which may or may not be biochemically meaningful, may also present additional challenges in processing the diffuse scattering data. Nevertheless, a small but growing number of systems have shown simpler patterns of strong diffuse features that appear to be connected to protein motions. Modeling of diffuse scattering for such systems has improved substantially in recent years. For some models, like normal modes, the agreement with the diffuse signal is still relatively weak (Fig. 2); for others, such as MD, the agreement is stronger [33], but still lower than what is typical in Bragg analysis.

What will be required to take diffuse scattering to the next level and make it an equal partner to Bragg analysis? We can look to traditional crystallography for clues. In Bragg analysis, using model phases, the details of the atomic structure only are revealed when the model of the whole system is sufficiently accurate. As the accuracy is increased, the effect of refining the model of small numbers of atoms can be seen in the analysis. When we have a sufficiently accurate model for all the diffracted X-rays, diffuse scattering might become sensitive to atomic details of correlated variations, with the caveat that the diffuse signal might contain less information than the Bragg diffraction, as measurements nearby in reciprocal space can be correlated. If atomic details of correlated motions can be revealed for the systems that have the clearest diffuse signal, diffuse scattering would then begin to provide very interesting information for these systems. As capabilities evolve, more complex patterns might be tackled and the analysis methods applied more generally. Meanwhile, it will be important to scour the diffraction image repositories such as SBGrid Databank [65] or CXIdb [66] and archive raw data from new experiments, identifying cases where the diffuse signal is strong and amenable to analysis [67].

Despite being present in all macromolecular diffraction patterns, the origins of diffuse scattering in protein crystallography are in many cases still mysterious. Our current short-term outlook is that, for a small number of cases, whether it is due to long-range [28] or short-range disorder [19,31,33], diffuse scattering will provide valuable information for structural modeling. The types of conformational heterogeneity that can be validated and, potentially, refined against diffuse scattering data can guide us to define better models of protein structure and dynamics. As the structural biology toolkit expands, X-ray scattering, including diffuse scattering, still provides unique capabilities to probe conformational ensembles over many length scales, as captured in the recent review by Meisburger et al. [26]. Ultimately, the better models of concerted motions will have far ranging impact beyond the average structure that is accessible using conventional X-ray crystallography and cryo-electron microscopy data, yielding a deeper understanding of biochemical mechanism [68].

## Acknowledgements

We thank TJ Lane, R Stroud, H Chapman, and K Ayyer, GN Phillips, Jr, and anonymous reviewers for helpful comments on the preprint. J.S.F. is supported by a Pew Scholar Award from the Pew Charitable Trusts, a Packard Fellowship from the David and Lucile Packard Foundation, NSF STC-1231306, NIH GM123159, and NIH GM124149. J.S.F. and M.E.W. are supported by UC Office of the President Laboratory Fees Research Program LFR-17-476732. M.E.W. is supported by US Department of Energy, via the Exascale Computing Project.

## Highlighted References

Polikanov YS, 2015

* Careful analysis of the diffuse scattering present in 70S ribosome crystals revealed that lattice vibrations may explain a significant portion of the diffuse signal, highlighting the importance of models that account for correlated variations across unit cell boundaries.

Van Benschoten AH, 2016

** Diffuse data is extracted from data collected under optimal conditions for Bragg analysis, revealing that modern PAD detectors and fine phi slicing can make diffuse data widely available. Also, LLM and normal modes models of disorder account for a substantial portion of the diffuse signal isolated from cypa and trypsin.

Ayyer K, 2016

** This work extends the analysis of diffuse X-ray scattering into the realm of XFELS and serial crystallography, while also advocating for the use of diffuse scattering for phasing and resolution extension. Rigid body translations of the PSII dimer are the proposed source of the diffuse signal.

Sui, S, 2016

* Graphene coated microfluidic chips enable collection of diffraction data with very high signal-to-noise, and may provide an alternative to jet based delivery systems for SFX experiments.

Peck A, 2017

** The authors approach diffuse scattering from a modelling perspective, and rigorously test current disorder models against a series of experimental datasets. Quantitative tests allow them to discern that most disorder models explain a limited portion of the diffuse signal, and that a LLM model is the best option, especially when correlations across unit cell boundaries are included.

Wall ME, 2017

*A molecular dynamics simulation of diffuse X-ray scattering from staphylococcal nuclease crystals is greatly improved when the unit cell model is expanded to a 2×2×2 layout of eight unit cells. The dynamics are dominated by internal protein motions rather than rigid packing interactions.

Van Benschoten AH, 2015

* Predicted diffuse scattering patterns differ substantially across different TLS models derived from the same data. This provides an important proof of principle for the use of diffuse scattering in refinement of macromolecular models.

Meinhold L, 2005

** In this study, the authors demonstrate the importance of long time-scale simulations to accurately sample a protein’s correlated motions. The improvement is most evident when examining the anisotropic portion of the diffuse signal attributed to protein dynamics alone.

Janowski PA, 2016

* MD simulations of lysozyme in a crystalline lattice reveal enhanced agreement with structural models derived from Bragg data. Nonetheless, convergence is slow, the lattice becomes disordered, and fluctuations of residues involved in crystal contacts are too high, indicating the need for improved MD force fields.

Meisburger, 2017

** This excellent review thoroughly lays out the connection between diffuse scattering, solution scattering, and crystallography. The assumptions and limitations of various approaches to analyzing diffuse data are clearly explained, and several disorder models are explored using case studies of biochemical interest.

